# Knock-in tagging in zebrafish facilitated by insertion into non-coding regions

**DOI:** 10.1101/2021.07.08.451679

**Authors:** Daniel S. Levic, Naoya Yamaguchi, Siyao Wang, Holger Knaut, Michel Bagnat

## Abstract

Zebrafish provide an excellent model for *in vivo* cell biology studies due to their amenability to live imaging. Protein visualization in zebrafish has traditionally relied on overexpression of fluorescently tagged proteins from heterologous promoters, making it difficult to recapitulate endogenous expression patterns and protein function. One way to circumvent this problem is to tag the proteins by modifying their endogenous genomic loci. Such an approach is not widely available to zebrafish researchers due to inefficient homologous recombination and the error-prone nature of targeted integration in zebrafish. Here, we report a simple approach for tagging proteins in zebrafish on their N- or C termini with fluorescent markers by inserting PCR-generated donor amplicons into non-coding regions of the corresponding genes. Using this approach, we generated endogenously tagged alleles for several genes critical for epithelial biology and organ development including the tight junction components ZO-1 and Cldn15la, the trafficking effector Rab11a, and the ECM receptor β1 integrin. Our approach facilitates the generation of knock-in lines in zebrafish, opening the way for accurate quantitative imaging studies.

**Summary statement:** Generation of endogenously tagged stable zebrafish knock-in lines is simplified by the integration of fluorescent protein cassettes with mRNA splicing elements into non-coding regions of genes.

## Introduction

Live imaging of fluorescently tagged proteins is an important tool for understanding gene function. In zebrafish, protein visualization has traditionally relied on overexpression studies in which tagged proteins are generated by microinjection of synthetic mRNA or by random integration of small transgenes (Kwan et al., 2007). These approaches are inherently limited because overexpression seldom recapitulates endogenous gene expression levels and spatiotemporal patterns. Large bacterial artificial chromosome (BAC) transgenes (Bussmann and Schulte-Merker, 2011, Navis et al., 2013, Alvers et al., 2014, Rodriguez-Fraticelli et al., 2015, Fuentes et al., 2016) can recapitulate the endogenous expression pattern and levels if they include the necessary regulatory sequences. However, regulatory sequences can reside hundreds of kilobases up- and downstream from the gene of interest and may be too far away to be included on a BAC transgene. Moreover, expression levels of BAC transgenes have been demonstrated to vary depending on the genomic insertion site (Fuentes et al., 2016) and the number of BAC transgene copies that are inserted (Chandler et al., 2007). The most accurate way to recapitulate physiological gene expression level and pattern is to tag the gene directly at its endogenous genomic locus to produce fusion proteins (Gibson et al., 2013), an approach commonly referred to as knock-in (KI). Recently, with the advent of CRISPR-Cas9 gene editing, KI has become feasible in cell lines and many model organisms (Dickinson et al., 2015, Koles et al., 2016, Dewari et al., 2018, Gao et al., 2019, Cronan and Tobin, 2019). In zebrafish, homology-based methods for generating KI lines have been described (Hoshijima et al., 2016, Wierson et al., 2020, Ranawakage et al., 2021). However, the available methods are difficult to implement and rely on inefficient DNA repair pathways, which are required for the precise integration of DNA the size of fluorescent protein sequences into the genome (Peng et al., 2014). As a result, only a few zebrafish KI lines expressing fluorescently tagged fusion proteins have been reported thus far. Moreover, endogenous N-terminal tagging, with the exception of the insertion of small peptides (Hoshijima et al., 2016, Ranawakage et al., 2021), has not been feasible in zebrafish. To circumvent these challenges, we devised a simple KI approach where precise integration is not needed for expression of endogenously tagged proteins. We targeted non-coding regions of genes, such as introns and 5’ untranslated regions (5’ UTRs), for integration of targeting cassettes that code for fluorescent proteins. In this manner, imprecise integration events do not affect the expression of the tagged proteins because non-coding sequences targeted by CRISPR-Cas9 are removed from the transcripts during RNA splicing. We used this approach to generate stable zebrafish KI lines for several proteins tagged on their N- or C-terminus by integrating the sequence for fluorescent proteins at the endogenous genomic loci. KI tagging in zebrafish opens the door for quantitative imaging studies, such as quantifying endogenous protein levels. As proof of principle, we measured the concentration of endogenous eGFP-Rab11a molecules on apical vesicles within different epithelial organs.

## Results and Discussion

To establish a method for endogenously tagging proteins in zebrafish, we devised an approach to integrate cassettes coding for fluorescent proteins into non-coding regions of genes. For C-terminal tagging, we used CRISPR-Cas9 to induce a double strand DNA (dsDNA) break in the intron preceding the last exon (Fig. 1A). We induced cutting in the intron at least 100 base pairs (bp) upstream of the last exon to minimize potential interference with pre-mRNA splicing (Fig. 1A). Together with Cas9 and the gRNA, we co-injected linear dsDNA PCR donor amplicons spanning part of the last intron and the coding sequence of the last exon of the gene of interest fused to the coding sequence of a fluorescent protein and a polyadenylation (polyA) sequence (Fig. 1B). The intron serves as a splice acceptor element. Integration of the donor can proceed through non-homologous end joining (NHEJ) at the 5’ and 3’ ends (Fig. 1C). Provided the cassette integrates in the correct orientation, expression of the modified transcript can tolerate errors, such as insertion-deletion mutations (INDELs), at integration boundaries because these sequences are removed during RNA splicing (Fig. 1D). Tagging the C-terminus of many proteins can impair their function. To circumvent this potential limitation, we also devised a strategy to add tags to the N-terminus of proteins. We induced dsDNA breaks in non-coding regions upstream of the start codon of the gene of interest using CRISPR-Cas9 (Fig, 1E). As described above, we co-injected a PCR donor amplicon, but the N-terminal tagging repair donors contain an approximately 500 bp homology arm (the endogenous sequence upstream from the dsDNA break). The homology arm was fused to the remaining upstream non-coding sequence, the coding sequence of a fluorescent protein, the coding sequence of the endogenous exon that harbors the start codon, and a portion of the following intron, which functions as a splice donor element (Fig. 1F). To promote homology-directed repair (HDR)-mediated integration at the 5’ end of the repair donor in the UTR, we mutated the gRNA target site in the repair donor (Fig. 1F). Integration of the 3’ end of the N-terminal repair donor does not require precise insertion by HDR because the intronic portion of the donor sequence is removed from transcripts during RNA splicing. (Fig. 1G-H).

**Figure 1.**
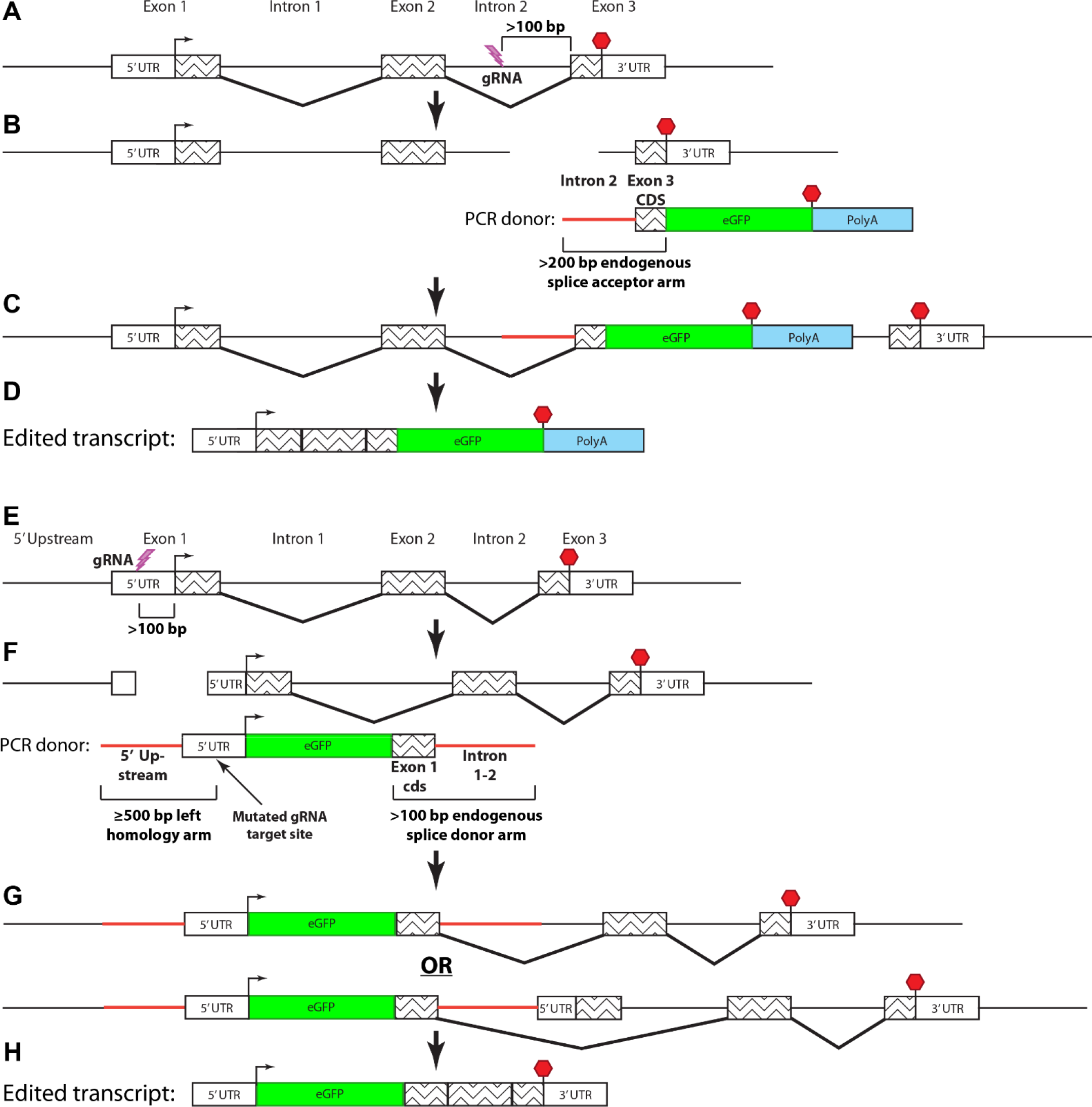
Knock-in fusion gene tagging in zebrafish using splice donor and acceptor arms. **(A-D)** C-terminal endogenous tagging strategy. **(A)** An intron upstream from the exon encoding the stop codon is targeted by CRISPR. **(B)** A dsDNA PCR donor containing a 5’ splice acceptor element (~200 bp of intron sequence), last exon coding sequence fused to a fluorescent protein coding sequence, and poly-adenylation sequence is co-injected. **(C-D)** Expression of the tagged transcript can tolerate error-prone integration of the PCR donor into the intron as it depends on mRNA splicing rather than precise genomic editing. **(E-H)** N-terminal endogenous tagging strategy. **(E)** A non-coding region upstream from the start codon is targeted by CRISPR. **(F)** A dsDNA containing a 5’ homology arm (~500 bp of sequence upstream from the gRNA target site), 5’ UTR with a mutated gRNA target site, fluorescent protein coding sequence, first exon coding sequence, and 3’ splice donor element (~100 bp of intron sequence) is co-injected. **(G-H)** Expression of the tagged protein occurs either if the 3’ end of the PCR donor is integrated by HDR (G, upper) or NHEJ (G, lower) because the endogenous first exon does not contain a splice acceptor site.

For certain endogenously tagged alleles, such as p2A-Cre or fusions to low-expressing genes, visual screening of a fluorescent protein is not feasible in the absence of other markers or transgenes. Therefore, we also devised a slightly modified approach for inserting larger sequences. We constructed a donor plasmid in which an endogenous splice acceptor element and the coding sequence of the last exon is cloned upstream of a tag of interest followed by a polyA sequence. The splice acceptor element in the donor plasmid is targeted and linearized by the same gRNA target site as the endogenous genomic intron. Integration of the donor and expression of the modified transcript follows similar principles as described for PCR donors, but this approach allows for the addition of larger cassettes that include transgenesis markers. Using this strategy, insertions can be identified based on the expression of the transgenesis marker but orientation of the insertion and expression of the tagged protein need to be confirmed by sequencing across the modified genomic region and by immunohistochemistry, respectively.

Using these approaches, we tagged several genes that have critical roles in epithelial biology and morphogenesis at their endogenous genomic loci. We injected a knock-in (KI) cocktail containing Cas9 protein, sgRNAs or synthetic crRNA/tracrRNA complexes, and PCR donor amplicons or plasmid into 1-cell stage embryos. We then visually screened embryos for fluorescence and compared it to the reported spatiotemporal mRNA expression patterns of the endogenous transcripts (Howe et al., 2021). We consistently observed expression of the fusion proteins in 1-5% of the injected embryos. Injected embryos with fluorescent protein expression showed varying levels of mosaicism (Fig. S1A-C). This mosaicism, combined with the highly restricted subcellular localization and endogenous levels of some proteins like aPKC (encoded by *prkci*) and ZO1(encoded by *tjp1a*), required us to use confocal microscopy to identify embryos with insertions of fluorescent protein encoding repair donor sequences (Fig. S1A-C).

To optimize conditions for endogenous tagging in zebrafish, we focused on *rab11a* because this gene exhibits widespread expression throughout embryonic and larval stages and insertion of the repair donor can easily be scored by visual screening (Fig. S2A). We prepared PCR donor amplicons for *rab11a* while modifying the gRNA target site position and mutating the gRNA sequence on the donor. We also compared dsDNA and single-stranded DNA (ssDNA) as repair donors. Although mutating the gRNA target site on the donor was not required for KI, this resulted in a >2-fold increase in efficiency (Fig. S2B-C). However, inducing dsDNA breaks on both the 5’ and 3’ ends of the integration site did not enhance KI efficiency, nor did using ssDNA as repair donor instead of dsDNA (Fig. S2B-C).

To determine the germline transmission efficiency of our approach, we targeted three genes (*cldn15la*, *rab11a*, and *tjp1a*) and raised only those injected embryos that showed mosaic expression to adulthood (F0). We then outcrossed 3–5 F0 animals for each target to wild-type (WT) fish and determined the percentage of stably expressing F1 embryos. *cldn15la* and *tjp1a*, targeted for C-terminal tagging of the gene products, showed similar levels of germline transmission rates (Fig. S3). Overall, targeting *cldn15la* and *tjp1a* resulted in 26.9% and 17.1% of F1 progeny showing expression, respectively (Fig. S3). Interestingly, we observed similar levels of efficiency despite using tdTomato (~1400 bp) as a tag for *cldn15la* and eGFP (~700 bp) for *tjp1a*, suggesting that insertion sizes in this range likely do not impact KI efficiency. For N-terminal tagging, we compared *rab11a* KI using dsDNA or ssDNA as donor repair sequences. Similar to F0 expression efficiency, using ssDNA as a repair donor did not improve the efficiency of obtaining stably expressing F1 embryos (Fig. S3), indicating that simple dsDNA PCR donor amplicons are effective for KI in zebrafish.

We next monitored the spatiotemporal expression patterns and subcellular localization of endogenously tagged proteins in stably expressing F2 larvae. ZO1 (Tjp1a) is a peripheral component of tight junctions and is localized cortically in epithelial cells (Zihni et al., 2016). ZO1-tdTomato showed widespread expression in epithelial organs through the body at embryonic and larval stages, with enrichment in the lens, floor plate, neural tube, vasculature, and intestine (Fig. 2A). In the eye, ZO1-tdTomato was present in the lens epithelium and was also enriched in closely apposed lens fiber cells (Fig. 2B, Movie 1). Close examination of ZO1-tdTomato in the trunk revealed enrichment within the dorsal aorta, caudal vein, intersegmental vessels, and notochord sheath cells (Fig. 2C, Movie 2). ZO1-tdTomato also labeled epithelial cells throughout the otic capsule, including sensory cells of cristae and those lining the canals (Fig. 2D, Movie 3). We also generated a line expressing endogenously tagged ZO1-eGFP and when crossed to the ZO1-tdTomato line, the two proteins showed identical expression patterns and co-localized intracellularly (Fig. S4A). In the epidermis, endogenously tagged ZO1 was highly expressed in lateral line neuromasts while in periderm cells it showed enrichment at tricellular junctions (Fig. S4B-C).

**Figure 2.**
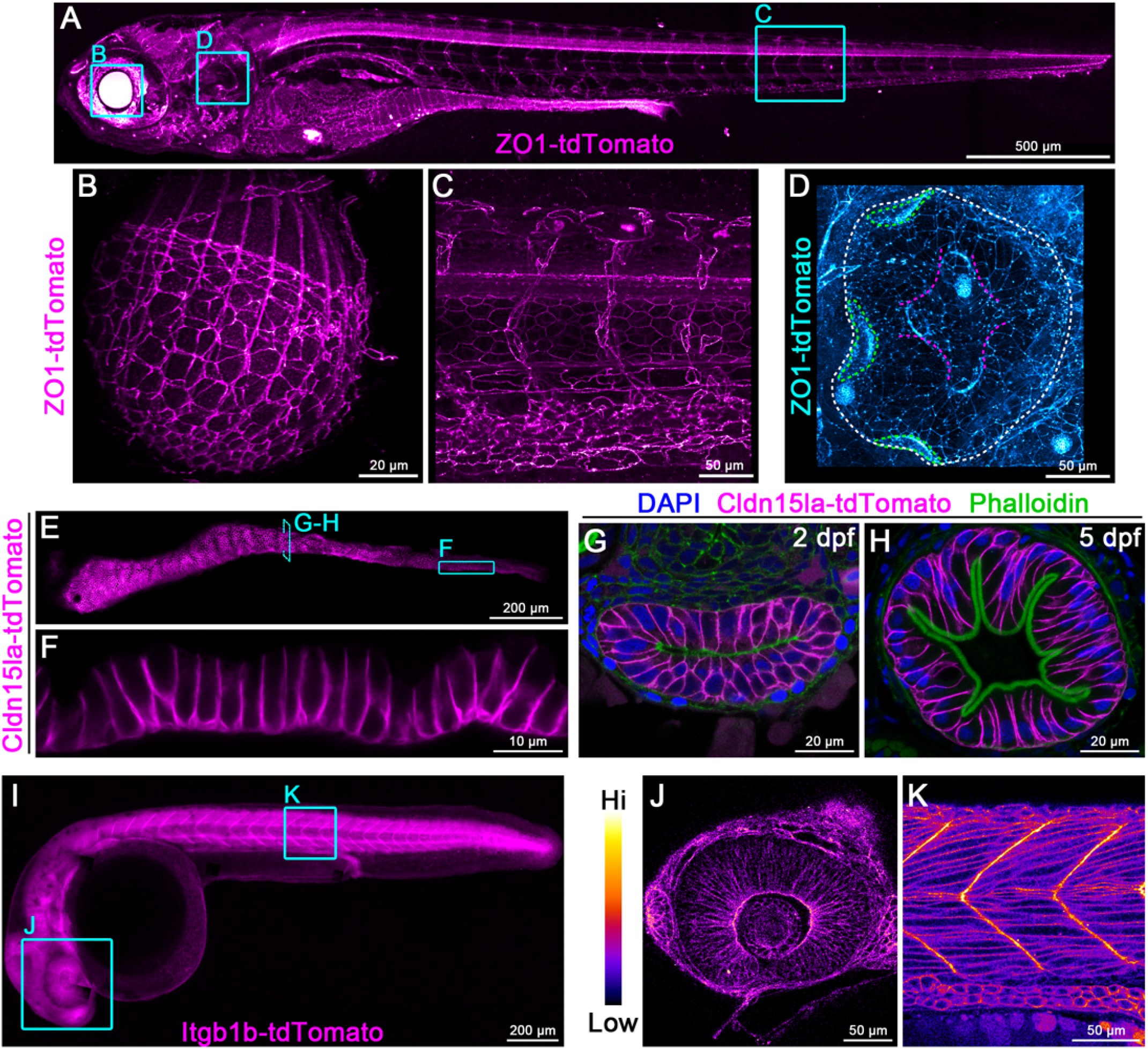
Endogenous C-terminal tagging of ZO1 (*tjp1a*) and Cldn15la (*cldn15la*). **(A-D)** Live imaging 3d reconstructions of *TgKI(tjp1a-tdTomato)*^pd1224^ heterozygous larvae. Cyan boxes in panel A show representative ROIs for panels B-D. For panel D: White dotted line, otic capsule; green dotted lines, cristae; magenta dotted lines, canals and septum. Panels A-C are pseudo-colored with the ImageJ/FIJI Magenta Hot LUT. Panel D is pseudo-colored with the ImageJ/FIJI Cyan Hot LUT. Animals are 7 dpf **(A)**, 5 dpf **(B)**, and 3 dpf **(C-D)**. Scale bars are 500 μm **(A)**, 20 μm **(B)**, and 50 μm **(C-D)**. **(E-F)** Live imaging of the intestine of a 7 dpf *TgKI(cldn15la-tdTomato)*^pd1249^ heterozygous larva. Panel E is pseudo-colored with the ImageJ/FIJI Magenta Hot LUT. Scale bars are 200 μm **(E)** and 10 μm **(F)**. **(G-H)** Transverse sections of the mid-intestine at the stages of lumen opening (2 dpf) **(G)** and onset of larval feeding (5 dpf) **(H)**. Scale bars are 20 μm. **(I-K)** Live imaging of a 28 hpf *TgKI(itgb1b-tdTomato)*^sk108^ heterozygous embryo. Cyan boxes in Panel I are representative ROIs for panels J-K, which are pseudo-colored according to the LUT scale shown. Scale bars are 200 μm **(I)** and 50 μm **(J-K)**.

To specifically label intestinal epithelial cells (IECs), we targeted *cldn15la*, which encodes a member of the claudin family of tetraspanin membrane proteins that regulate tight junction assembly and function. Cldn15la is an atypical member of the claudin family that is not restricted to tight junctions and is instead localized along basolateral membranes (Alvers et al., 2014). Similar to the transgenic *TgBAC(cldn15la-GFP)*^*pd1034*^ allele (Alvers et al., 2014), endogenously tagged Cldn15la-tdTomato was expressed in all IECs (Fig. 2E-F), where it was present on basolateral membranes throughout gut development (Fig. 2G-H). Of note, unlike the *TgBAC(cldn15la-GFP)*^*pd1034*^ allele which is homozygous lethal, *TgKI(cldn15la-tdTomato)*^*pd1249*^ was well tolerated in homozygous animals.

To visualize cell-ECM adhesions with an endogenous protein in live zebrafish, we generated an *itgb1b-tdTomato* KI line. Consistent with prior studies (Martinez-Morales et al., 2009, Sidhaye and Norden, 2017), Itgb1b-tdTomato was clearly enriched at the basal membrane of the cells in the optic cup at 28 hpf. We also confirmed prominent enrichment of Itgb1b-tdTomato in myotendinous junctions at the somite boundaries (JüLich et al., 2005). These observations suggest that *itgb1b-tdTomato* faithfully reports the localization of Itgb1b in live zebrafish.

To establish a zebrafish model for *in vivo* studies of membrane trafficking, we generated a KI line expressing N-terminally tagged eGFP-Rab11a. Live imaging revealed that eGFP-Rab11a is nearly ubiquitously expressed and enriched in epithelial organs (Fig. 3A, Movie 4). In transverse sections of the posterior intestine, eGFP-Rab11a was highly expressed and localized apically in lysosome-rich enterocytes (LREs) (Park et al., 2019) and epithelial cells of the pronephros (Fig. 3B). The apical localization of eGFP-Rab11a in zebrafish IECs resembles that of endogenous Rab11a by immunostaining (Levic et al., 2020). Among lower expressing tissues, eGFP-Rab11a was still easily detected in transverse sections of skeletal muscle, notochord sheath cells, and notochord vacuolated cells (Fig. 3C). Live imaging revealed dynamic movement of eGFP-Rab11a in the cytoplasm of notochord vacuolated cells (Movie 5). We also noted high enrichment of eGFP-Rab11a in lateral line neuromasts by live imaging (Fig. 3A). Within neuromasts, eGFP-Rab11a expression was restricted to hair cells, where it was enriched apically near stereocilia of the apical membrane (Fig. 3D). eGFP-Rab11a was also present at basal puncta that may represent contact sites from neurons that innervate neuromasts (Fig. 3D). Accordingly, we detected enriched expression of eGFP-Rab11a in tracts and projections of neurons that underlie neuromasts (Fig. 3E-F).

**Figure 3.**
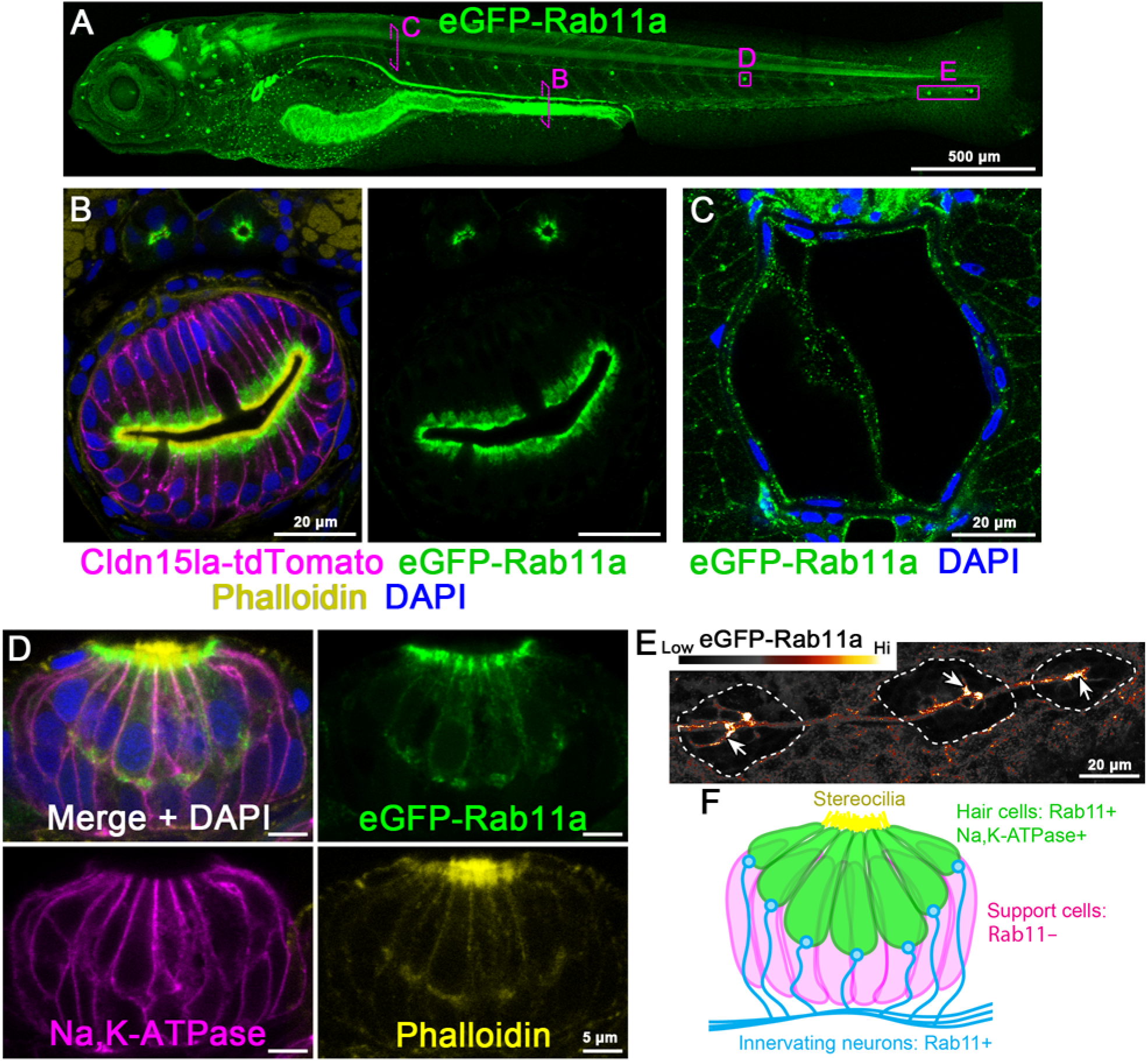
Endogenous N-terminal tagging of Rab11a (*rab11a*). **(A)** Live imaging 3d reconstruction of 5 dpf *TgKI(eGFP-rab11a)*^pd1244^ heterozygous larva. Magenta boxes show representative ROIs for panels B-E. Scale bars is 500 μm. **(B)** Transverse section through the posterior mid-intestine (LREs) and pronephric ducts. Scale bars are 20 μm. **(C)** Transverse section through the notochord. Scale bars is 20 μm. **(D)** Whole mount image of a neuromast. Scale bars are 5 μm. **(E)** Live lateral image of neurons innervating the terminal lateral line neuromasts (outlined by dotted line). Image is pseudo-colored according to the LUT scale shown. Scale bars is 20 μm. **(F)** Schematic summarizing expression and localization of Rab11a within neuromasts.

Having shown that proteins can be tagged endogenously we tested whether our KI lines can be used to determine their relative cellular abundance. To explore this question, we adapted a microscopy-based approach to measure the concentration of GFP molecules on diffraction limited vesicles on zebrafish tissue sections to estimate the number of eGFP-Rab11a molecules per vesicle. While this technique is commonly used in *in vitro* models (Clayton, 2018, Marques et al., 2019, Escamilla-Ayala et al., 2020), quantitative imaging of animal tissues can be obscured by factors such as autofluorescence and light scattering. Therefore, to establish baseline standards, we measured the photon emission of purified eGFP particles when imaged on zebrafish tissue sections. We collected intestinal tissue sections of eGFP-negative WT larvae (Fig. 4A), incubated a solution of purified eGFP at low concentration, and then crosslinked eGFP particles to the tissue surface by fixation (Fig. 4B). We collected accumulated photon counts of eGFP particles and then photobleached them (Fig. 4C). During photobleaching we monitored signal intensity and inferred the original number of eGFP molecules in the particle based on the signal decay profile (Fig. 4D). Using this approach, we identified eGFP particles containing 1-3 molecules that exhibited a linear increase in photon emission (Fig. 4E-F). Coincidentally, background level emission from tissue autofluorescence generated approximately the same number of photon counts as a single eGFP molecule, on average (Fig. 4F). Next, we prepared eGFP-Rab11a KI larvae and performed single particle imaging of diffraction limited apical vesicles on tissue sections (Fig. 4G-H) using identical processing and imaging conditions as described above. Although Rab11a expression levels vary by more than 2-fold at the mRNA levels in LREs vs. IECs (Park et al., 2019), endogenously tagged eGFP-Rab11a concentration on apical vesicles did not change in proportion (Fig. 4I). However, the relative distribution profile for LREs did reveal an increase in the fraction of vesicles containing >3 molecules (Fig. 4J), possibly reflecting a pool of vesicles that function in protein uptake in these specialized enterocytes (Park et al., 2019).

**Figure 4.**
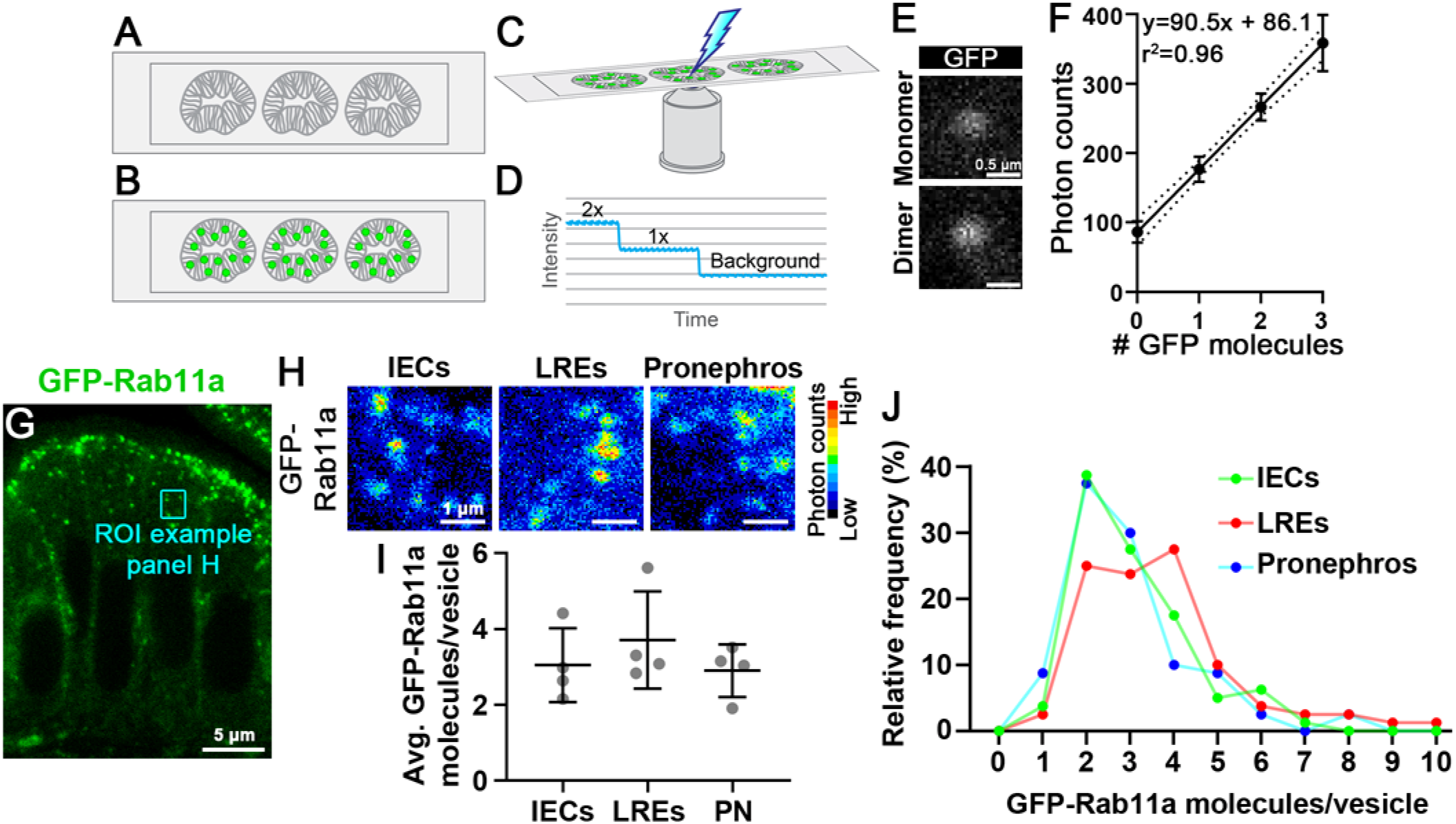
Tissue-specific Rab11a expression levels do not strongly affect its concentration on apical vesicles of the intestine or pronephros. **(A-D)** Schematic outlining single particle imaging to measure photon emission of purified GFP on zebrafish tissue. Intestinal sections of GFP-negative larvae are collected **(A)** and GFP solution (340 ng/mL) is incubated with and crosslinked onto the tissue surface **(B)**. Photon counts of GFP particles are collected and the particles are photobleached to background-level intensity **(C)**. The number of GFP molecules per particle are inferred by the decay profile. **(E-F)** Single particle photon count imaging **(E)** and linear regression analysis of purified GFP photon emission when imaged on intestinal tissue sections. Scale bars are 0.5 μm. **(G)** Transverse section of mid-intestine enterocytes of a *TgKI(eGFP-rab11a)*^pd1244^ heterozygous larva. Cyan box shows a representative ROI of a group of apical vesicles used for single particle imaging. **(H)** Pseudo-colored photon count images of apical vesicles of mid-intestine enterocytes (IECs), posterior mid-intestine lysosome-rich enterocytes (LREs), and pronephric duct epithelial cells (PN). Photon count intensity LUT scale is shown on the right. Scale bars are 1 μm. **(I)** Plot of average GFP-Rab11a concentration values from apical vesicles. Data points are average values from tissue sections of individual larvae. n=4 larvae for each organ (20 vesicles per animal). Data were not significantly different (One-way ANOVA). **(J)** Relative frequency plot of the data used for panel I. LREs vs. IECs, p<0.05; LREs vs. PN, p<0.01; IECs vs. PN, not significant (One-way ANOVA). n=80 vesicles per organ.

In this study, we describe a simple and effective approach to tag genes with targeting cassettes encoding fluorescent proteins at their endogenous genomic loci in zebrafish. While CRISPR and Zinc Finger Nuclease gene editing have been used to generate knock-in zebrafish lines to study promoter activity (Kimura et al., 2014, Hoshijima et al., 2016, Li et al., 2019), there are few published examples of C-terminally tagged KI fusion lines (Cronan and Tobin, 2019) and, with the exception of small peptide insertions (Hoshijima et al., 2016, Ranawakage et al., 2021), there are no reported examples of stable zebrafish KI lines of N-terminally tagged proteins. This scarcity may reflect the highly error prone nature of HDR in zebrafish. Our approach is unique because integration errors, such as INDELs, in non-coding gene regions can still mediate proper expression at the protein level because integration boundaries are excluded by RNA splicing. During the preparation of this manuscript, a similar endogenous tagging approach was reported for cultured mammalian cells (Zhong et al., 2021). Additionally, a related technique for targeted protein trapping by intronic insertion of artificial exons has been demonstrated to work for cultured mammalian cells (Serebrenik et al., 2019). In contrast to these examples, we sought to minimize the integration of undesired plasmid DNA elements by injecting PCR donor amplicons that only encode the relevant functional elements. Following this approach, we generated stable zebrafish KI fusion lines for several integral membrane and membrane associated proteins critical for epithelial development and cell physiology. Using quantitative imaging, we used one of these stable lines to measure the concentration of eGFP-Rab11a molecules on apical vesicles in different epithelial organs. Importantly, because genes are tagged at the endogenous locus, the zebrafish lines presented here improve accuracy and allow experimental approaches not feasible with traditional transgenic lines. These include the abilities to recapitulate spatiotemporal expression patterns of endogenous genes and to precisely quantify protein levels using single particle imaging or related techniques like fluorescence correlation spectroscopy (Wang et al., 2018). They also can facilitate uncovering protein interaction networks and dynamics without overexpression artifacts (Ahmed et al., 2018) and acute manipulation of protein function using conditional loss-of-function approaches such as degron-mediated protein depletion (Daniel et al., 2018, Yamaguchi et al., 2019). One factor that can impact expression levels of endogenously tagged proteins is the 3’ UTR used in the repair donor sequence. The 3’ tagged lines shown here have exogenous polyA sequences that may alter stability of the modified transcript. If known, the endogenous 3’ UTR and polyA sequence can be substituted in the 3’ repair donor to recapitulate mRNA stability levels more accurately. By contrast, genes tagged with 5’ insertions, such as *eGFP-rab11a*, will result in transcripts containing the endogenous 3’UTR that can more closely provide endogenous expression levels. Finally, because our KI approach relies on splice donor and acceptor elements, our method can be adapted to generate internal insertions at precise codon positions for targets such as membrane proteins containing signal peptide sequences that cannot be tagged at their C-termini.

## Materials and methods

### Zebrafish maintenance

Zebrafish (*Danio rerio*) were used in accordance with Duke University Institutional Animal Care and Use Committee (IACUC) guidelines and NYU School of Medicine under the approval from protocol number 170105–02. Zebrafish stocks were maintained and bred as previously described (Westerfield, 2007). Genotypes were determined by PCR and DNA sequencing or phenotypic analysis. Male and female breeders from 3–18 months of age were used to generate fish for all experiments. 1–7 dpf zebrafish larvae from the Ekkwill (EK) or AB/TL background were used in this study. Strains generated in this study are: *TgKI(tjp1a-tdTomato)*^pd1224^, *TgKI(tjp1a-eGFP)*^pd1252^, *TgKI(cldn15la-tdTomato)*^pd1249^, *TgKI(itgb1b-tdTomato)*^sk108^, *TgKI(eGFP-rab11a)*^pd1244^. Embryos and larvae were anesthetized with 0.4 mg/ml MS-222 (Sigma, A5040) dissolved in embryo media for handling when necessary.

### Generation of C-terminal PCR donors

We first generated a series of donor vectors to expedite production of C-terminal knock-in constructs. In the pUC19 vector backbone, we constructed a multiple cloning site, a fluorescent protein coding sequence or other tag lacking the start codon (eGFP, mLanYFP, mScarlet, tdTomato, p2A-QF2, p2A-eGFP, p2A-mScarlet, or p2A-Venus-PEST), a stop codon, a second multiple cloning site, and the zebrafish *ubb* poly-adenylation sequence. This fragment was flanked by forward and revers PCR primer sites to generate PCR donor amplicons for all targets using the same primers, pUC19_forward 5’-GCGATTAAGTTGGGTAACGC-3’ and pUC19_reverse 5’-TCCGGCTCGTATGTTGTGTG-3’. A gene fragment spanning from the middle of the last intron through the last coding sequence codon of the exon was cloned into the donor vector in frame with the fluorescent protein coding sequence using the following primers: tjp1a-forward 5’-cttgctagcAGTTTCGATGACCACAGGGT-3’, tjp1a-reverse 5’-cctctcgagGAAATGGTCAATAAGCACAGACA-3’, cldn15la-forward 5’-cttccgcggGTTTCACGTCAGAAATTGTCGG-3’, cldn15la-reverse 5’-cttctcgagGACGTAGGCTTTGGATGTTTC-3’. PCR donor amplicons were purified using the Nucleospin Gel and PCR Clean-up kit (Machery-Nagel, distributed by Takara Bio USA). PCR products were not gel purified. The final product was dried on column at 60°C for 10 minutes and then eluted with water and stored at −20°C. C-terminal donor vectors will be deposited to Addgene.

### Generation of N-terminal PCR donors for *rab11a*

A gene fragment spanning from 446 base pairs upstream from the 5’ UTR through 491 base pairs downstream of the end of exon 1 was cloned into pCS2+ with the following primers: rab11a-forward 5’-cttctcgagGAACTTACGAGCTGGATTTGTGC-3’ and rab11a-reverse 5-ctttctagaTGACAGCGTCGGTCACAGTT-3’. A small multiple cloning site was added before the start codon of exon 1 by site directed mutagenesis (Q5 SDM Kit, New England Biolabs) using the primers rab11a-MCS-SDM-forward 5’-tactagttccATGGGGACACGAGACGAC-3’ and rab11a-MCS-SDM-reverse 5’-agaccggtaggCTCGATCAAAACAAAAGCGC-3’. eGFP was cloned into the multiple cloning site using the primers GFP-forward 5’-cttaccggtgccgccaccATGGTGAGCAAGGGCGAGGA-3’ and GFP-reverse 5’-cttactagtCTTGTACAGCTCGTCCATGCC-3’. The gRNA target sites used for genomic targeting were mutated in the donor plasmid using site directed mutagenesis with the primers: gRNA-1-SDM-forward 5’-aacagcgaactGTCGCCTCCACTTTCCTT-3’, gRNA-1-SDM-reverse 5’-atctccgctgtaGCACTGCAGTCTGTCTGT-3’, gRNA-2-SDM-forward 5’-actcgagcagagCAAACAAACTCCTGCTCTTC-3’, gRNA-2-SDM-reverse 5’-cgagctagcataTTAGCTGGCCTTTACTGT-3’. PCR donors were generated as described above using the primers: rab11a-donor-forward 5’-GAACTTACGAGCTGGATTTGTGC-3’ and rab11a-donor-reverse 5-ctttctagaTGACAGCGTCGGTCACAGTT-3’. For ssDNA production in Fig. S2, PCR of the same donor plasmid was performed using the same primers, but the forward primer was phosphorylated. After PCR ssDNA was generated using the Guide-it Long ssDNA Production System (Takara Bio USA) using the manufacturer’s recommendations. ssDNA was purified using the Nucleospin Gel and PCR Clean-up kit (Machery-Nagel, distributed by Takara Bio USA) with buffer NTC used as recommended by the manufacturer. ssDNA conversion was verified used gel electrophoresis and the product was stored at −80°C.

### Generation of N-terminal PCR donors for *prkci*

A gene fragment spanning from 488 base pairs upstream from the 5’UTR through 61 base pairs downstream from exon 1 was cloned into pDONR221 using a BP reaction (ThermoFisher) using the primers prkci-BP-forward 5’-GGGGACAAGTTTGTACAAAAAAGCAGGCTCctatctaggtatatgggccctc-3’ and prkci-BP-reverse 5’-GGGGACCACTTTGTACAAGAAAGCTGGGTcgcaatcctgagaataagtgaga-3’. The gRNA target site in intron 1 was mutated by PCR during initial cloning (the reverse cloning primer was mutagenic), and the gRNA target site sequence was verified independently in a population of WT fish. Next a multiple cloning site was inserted before the start codon of exon 1 using the primers prkci-MCS-SDM-forward 5’-tactagttccATGCCCACGCTGCGGGAC-3’ and prkci-MCS-SDM-reverse 5’-agaccggtaggTATGGACTATCCGTACTCCTGCTAGC-3’. eGFP was cloned to the site using the primers GFP-forward 5’-cttaccggtgccgccaccATGGTGAGCAAGGGCGAGG-3’ and GFP-reverse 5’-cttactagtCTTGTACAGCTCGTCCATGCC-3’. PCR donors were generated as described above using the primers prkci-forward-donor 5’-tatctaggtatatgggccctc-3’ and prkci-reverse-donor 5’-gcaatagtgcgaataagtgaga-3’.

### Generation of plasmid donors for *itgb1b*

A genomic fragment spanning the last 29 bp of itgb1b exon 8 to the end of *itgb1b* exon 9 was cloned into the pUC19 plasmid (exon numbering is based on transcript ID: ENSDART00000161711.2). A linker sequence coding for amino acids GGPVAT was inserted after the codon for the last amino acid of *itgb1b* and the fragment was fused to the tdTomato coding sequence followed by the SV40 polyA signal sequence. A gRNA target site was designed to target both the donor plasmid, thereby linearizing it in the intron, and the endogenous genomic intron.

### Production of guide RNA (gRNA)

Guide RNA (gRNA) target sites were identified using CRISPRscan (Moreno-Mateos et al., 2015) and gRNAs were synthesized by *in vitro* transcription using an oligo-based template method (Yin et al., 2015) using the MEGAshortscript T7 Transcription Kit (ThermoFisher). gRNAs were precipitated by ammonium acetate/isopropanol, resuspended in water, and stored at −80°C. For *itgb1b*, crRNA and the tracrRNA were purchased from IDT, and the cRNA was designed using the Custom Alt-R CRISPR-Cas9 guide RNA Design Tool (IDT). gRNA target sites used in this study were: *cldn15la*, 5’-GtTTCACGTCAGAAATTGTCGGG-3’ and 5’-GGATTTCTCTAGATTATGACCGG-3’; *prkci*, 5’-GcATTCTCACTTATTCTCAACGG-3’; *rab11a,* 5’-gGCAGCGGAGAGGACAGCGACGG-3’ and 5’-CCGGCTAGCTCACTTCGAGCAcC-3’; *tjp1a*, 5’-tGCGAATAGGGGTTGATAATGGG-3’ and 5’-GaGTTTCGATGACCACAGGGTGG-3’; crRNA for *itgb1b*, 5’-GGAGGTCTTGATGTAGGATT-3’.

### Microinjections and visual screening

Early 1-cell stage embryos were injected with 1-2 nL of a knock-in cocktail consisting of gRNA (final concentration 30-50 pg/nL), dsDNA or ssDNA PCR donors (final concentration 5-10 pg/nL) Cas9 protein tagged with a nuclear localization sequence (PNA Bio CP-01) (final concentration 300-500 pg/nL), and phenol red (final concentration 0.05%). We observed mortality rates of 10-20% for dsDNA-injected embryos and 40-50% for ssDNA-injected embryos. For *itgb1b*, the injection mix containing Cas9-NLS protein, crRNA, tracrRNA and the plasmid harboring the itgb1b targeting cassette was heat-activated at 37°C and injected into one-cell stage wild-type embryos. Embryos were visually screened daily between 1–5 dpf for fluorescence using an Axio Zoom V16 microscope (Zeiss). Embryos suspected of showing fluorescence were mounted in 0.7% low melting point agarose and imaged by confocal microscopy on a Leica SP8 microscope using an HC FLUOTAR VISIR 25x/0.95 NA water immersion objective (Leica). Positive embryos were recovered from anesthesia and raised to adulthood.

### Isolation of stable alleles

Injected embryos showing expression of fluorescently tagged proteins were raised and crossed to WT fish. The positive F1 embryos were raised and crossed to WT fish. The integration site was sequenced at the F2 generation and the lines were designated allele numbers. With the exception of Fig. S1-S2, all imaging data presented are from animals of the F2 or greater generation

### Imaging and image processing

All imaging was performed on a Leica SP8 confocal microscope. Live imaging was conducted with a FLUOTAR VISIR 25x/0.95 NA or HC PL APO CS2 20x/0.75 water immersion objectives (Leica), and cross sections with an HC PL APO CS2 63x/1.40 oil immersion objective (Leica). Whole animals were imaged in tiling mode and the data were stitched in Leica LAS software. Imaging data were processed in ImageJ/FIJI (NIH) to prepare 3D reconstructions using native plugins. To enhance visualization of some images, data were pseudo-colored using default lookup tables (LUTs) in ImageJ/FIJI. LUT scales are presented in the figure panels. Post-processing for linear changes in brightness were performed in photoshop using the levels tool.

### Single particle imaging

6 dpf GFP-negative larvae were fixed in 4% paraformaldehyde in PBS pH 7.5 overnight at 4°C, rinsed in PBS, and then embedded in 5% low melting point agarose. 200 μm sections were collected using a Leica VT1000S vibratome (Levic et al., 2020). Sections were incubated with 340 ng/mL purified eGFP, which was generated as we previously described (Park et al., 2019), overnight at 4°C. The solution was then gently aspirated and then sections were fixed in 4% paraformaldehyde in PBS pH 7.5 for 30 minutes at room temperature. Sections were rinsed in PBS and then mounted on glass slides in 90% glycerol buffered with 10 mM Tris, pH 8 with 1% N propyl-gallate added. Sections were imaged near the coverslip surface with a Leica SP8 confocal microscope using an HC PL APO CS2 63x/1.40 oil immersion objective. Excitation was performed with a 20 mW 488 nm laser operating at 0.2% power, and scans were performed at 400 Hz with a pixel size of 50 nm. Emission spectra were collected from 498-550 nm using a HyD detector operating in photon counting mode with 10x line accumulation and at 10% gain. Experimental samples (eGFP-Rab11a larvae) were processed identically. Raw 12-bit images were analyzed in ImageJ/FIJI. Photon counts of 5 pixel^2^ ROIs of eGFP particles were collected and analyzed by linear regression using Graphpad Prism. Photon counts experimental samples were interpolated from the linear regression analysis of purified eGFP particles to infer the number of eGFP molecules per vesicle.

## Acknowledgements

We thank the Duke Zebrafish Core and Joseph Proietti and Sam Pirani for excellent fish care and maintenance and Jieun Esther Park for providing purified eGFP. We also thank Xiaolei Wang, Ian Macara, and Kristen Kwan for helpful discussions. The use of the NYULH DART Microscopy Laboratory (P30CA016087) is gratefully acknowledged.

## Author contributions

D.S.L. developed the methodology for N-terminal endogenous tagging and C-terminal endogenous tagging, generated the *cldn15la*, *tjp1a*, and *rab11a* zebrafish lines, and collected imaging data. N.Y. developed the methodology for C-terminal tagging using a plasmid as a repair donor, generated the *itgb1b* zebrafish line, and collected imaging data. S.W. collected imaging data. D.S.L. wrote the manuscript with input from all authors. M.B. and H.K. supervised the project.

## Data availability

C-terminal knock-in donor vectors will be deposited to Addgene and stable knock-in zebrafish lines to ZIRC, but in the interim they are available from our labs upon request to the corresponding author. A detailed protocol of our endogenous tagging method is provided as a supplementary file. Sequence files describing the genomic integration sites for all KI alleles are provided as supplementary files.

---

Movie 1. **Rotating 3D reconstruction of ZO1-tdTomato expression in the lens**. Data are from Fig. 2B.

Movie 2. **Rotating 3D reconstruction of ZO1-tdTomato expression in the embryonic trunk**. Data are from Fig. 2C.

Movie 3. **Rotating 3D reconstruction of ZO1-tdTomato expression in the otic capsule**. Data are from Fig. 2D.

Movie 4. **Rotating 3D reconstruction eGFP-Rab11a expression in 5 dpf whole larvae**. Data are from Fig. 3A.

Movie 5. **Live imaging of eGFP-Rab11a vesicle dynamics in notochord vacuole cells**. Data are related to Fig. 3C. Data were acquired at 1 frame every 3 seconds.

Supplementary file 1. Sequence file for the genomic integration site of *TgKI(tjp1a-tdTomato)*^*pd1224*^.

Supplementary file 2. Sequence file for the genomic integration site of *TgKI(tjp1a-eGFP)*^*pd1252*^.

Supplementary file 3. Sequence file for the genomic integration site of *TgKI(cldn15la-tdTomato)*^*pd1249*^.

Supplementary file 4. Sequence file for the genomic integration site of *TgKI(itgb1b-tdTomato)*^*sk108*^.

Supplementary file 5. Sequence file for the genomic integration site of *TgKI(eGFP-rab11a)*^*pd1244*^.

**Figure S1.**
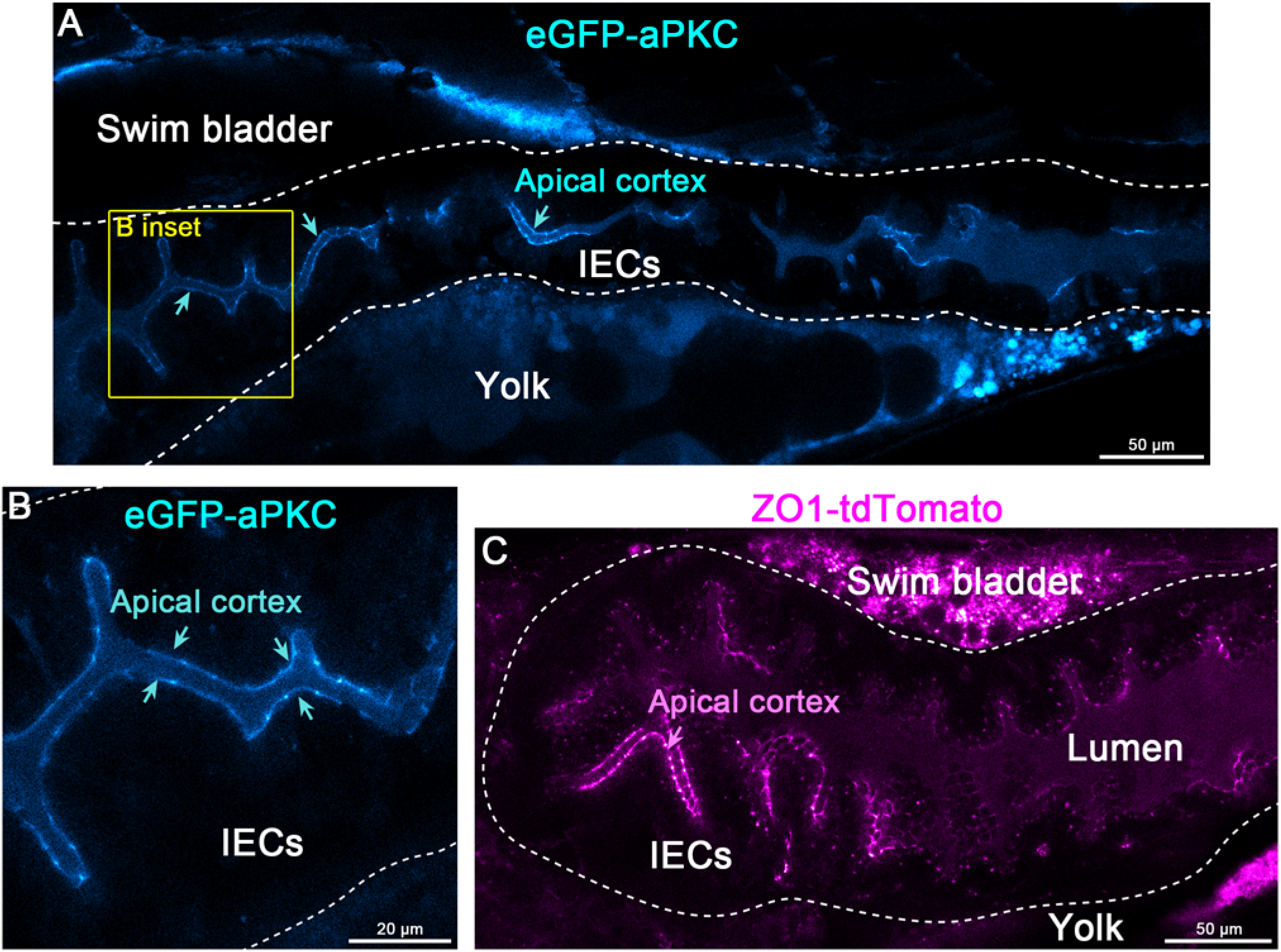
Examples of mosaicism in injected embryos (F0s). **(A-B)** Live imaging of a 5 dpf eGFP-aPKC (encoded by *prkci/has*) F0 larva. Cyan arrows point to the apical cortex. Panels A-B are pseudo-colored with the ImageJ/FIJI Cyan Hot LUT. Scale bars are 50 μm **(A)** and 20 μm **(B)**. **(C)** Live imaging of a 5 dpf ZO1-tdTomato (encoded by *tjp1a*) F0 larva. Panel C is pseudo-colored with the ImageJ/FIJI Magenta Hot LUT. Magenta arrow points to apical the cortex. Scale bars is 50 μm. The dotted line marks the intestinal epithelium.

**Figure S2.**
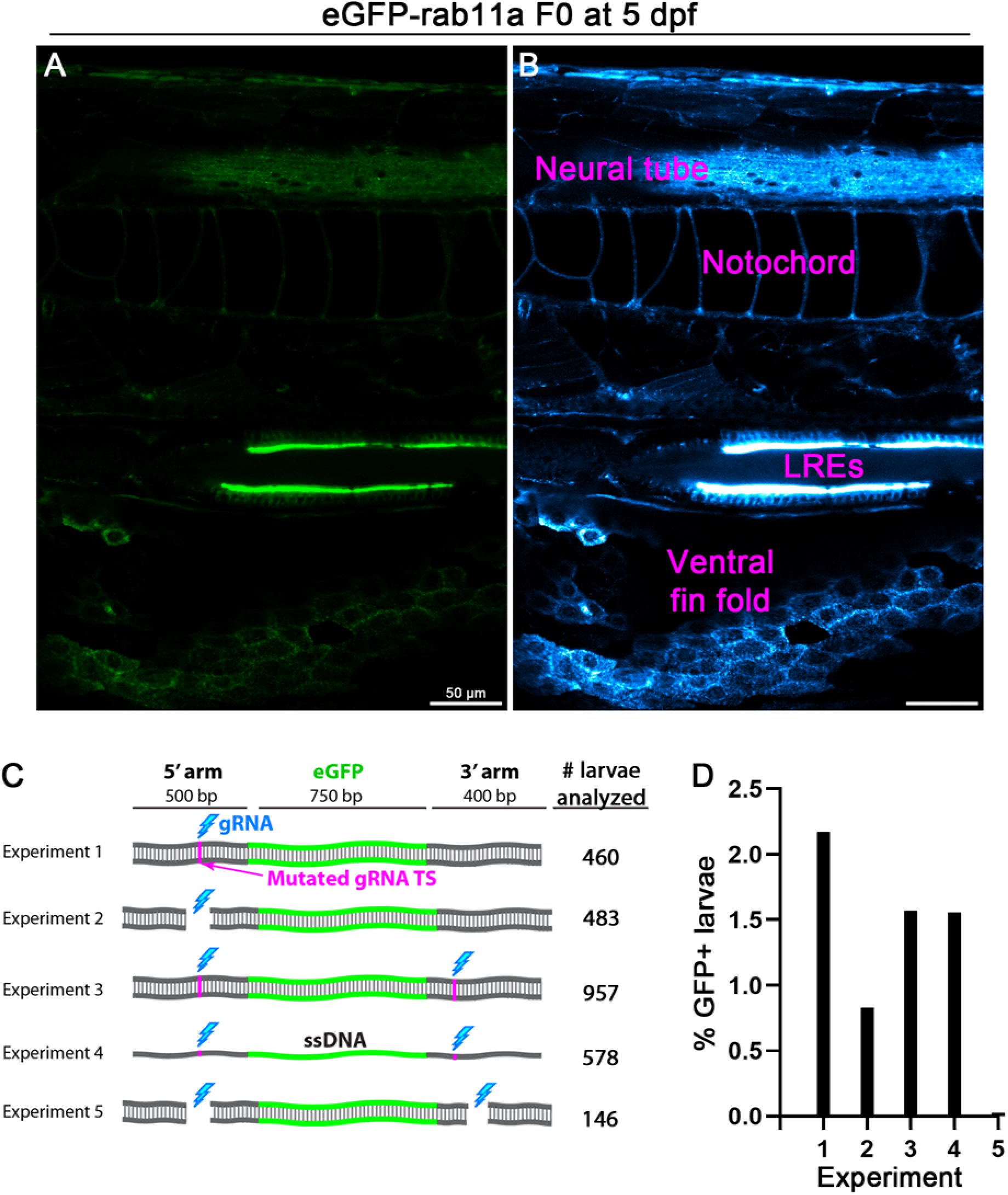
Variables affecting F0 efficiency for N-terminal endogenous tagging. **(A-B)** Representative examples of mosaic expression of eGFP-Rab11a in injected larvae (F0s). The broad spatial expression patterns and high expression levels of eGFP-Rab11a allow for simple visual screening to test PCR donor variables to optimize endogenous tagging. Panel A is original confocal live image, and panel B is pseudo-colored with the ImageJ/FIJI Cyan Hot LUT to enhance visualization of lower expressing cell-types. Scale bars are 50 μm. **(C)** 1-cell stage embryos were injected with a knock-in cocktail containing Cas9 protein, while the PCR donor and gRNAs varied. The cyan bolt indicates the relative position of the gRNA (upstream or downstream from the insertion site or both). The magenta line indicates when the PAM site for the gRNA was mutated on the PCR donor. For experiment 4, linear single-stranded DNA (ssDNA) was used rather than dsDNA PCR donors. The numbers to the right indicate the total number of surviving larvae that were visually screened for fluorescence. **(D)** Plot of F0 expression efficiency rates for the experiments in panel C.

**Figure S3.**
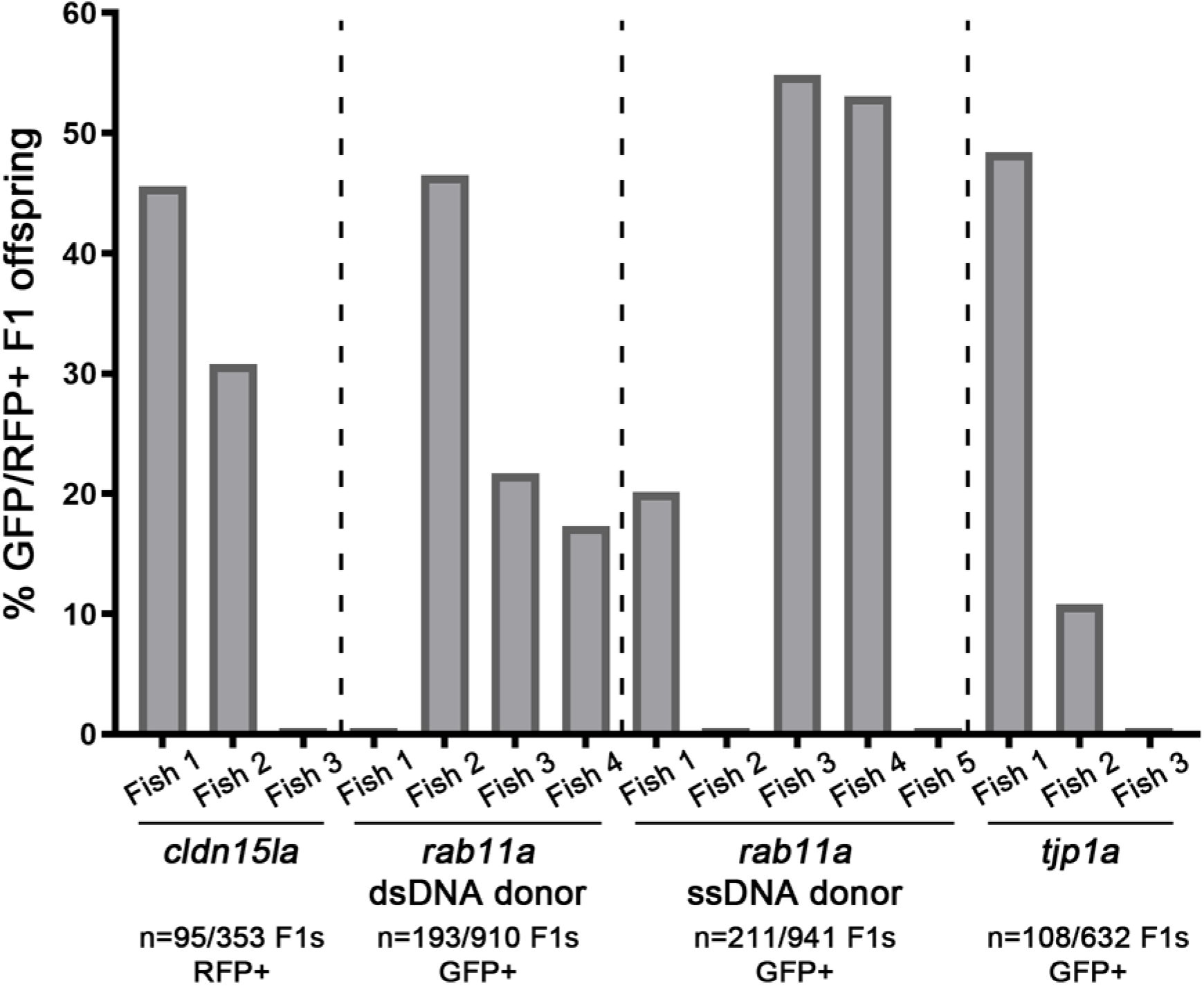
Germline transmission efficiency rates following visual screening of injected larvae. 1-cell stage embryos were injected with knock-in cocktails for different genes and then visually screened for fluorescence. The larvae showing expression were raised to maturity and then outcrossed. 3-5 animals were outcrossed per condition, and the F1 progeny were visually screened for fluorescence. The germlines of F0 adults have mosaic integration, so the knock-in alleles do not show Mendelian segregation in the F1 embryos. The data plotted indicates the levels of mosaicism present in the F0 generation.

**Figure S4.**
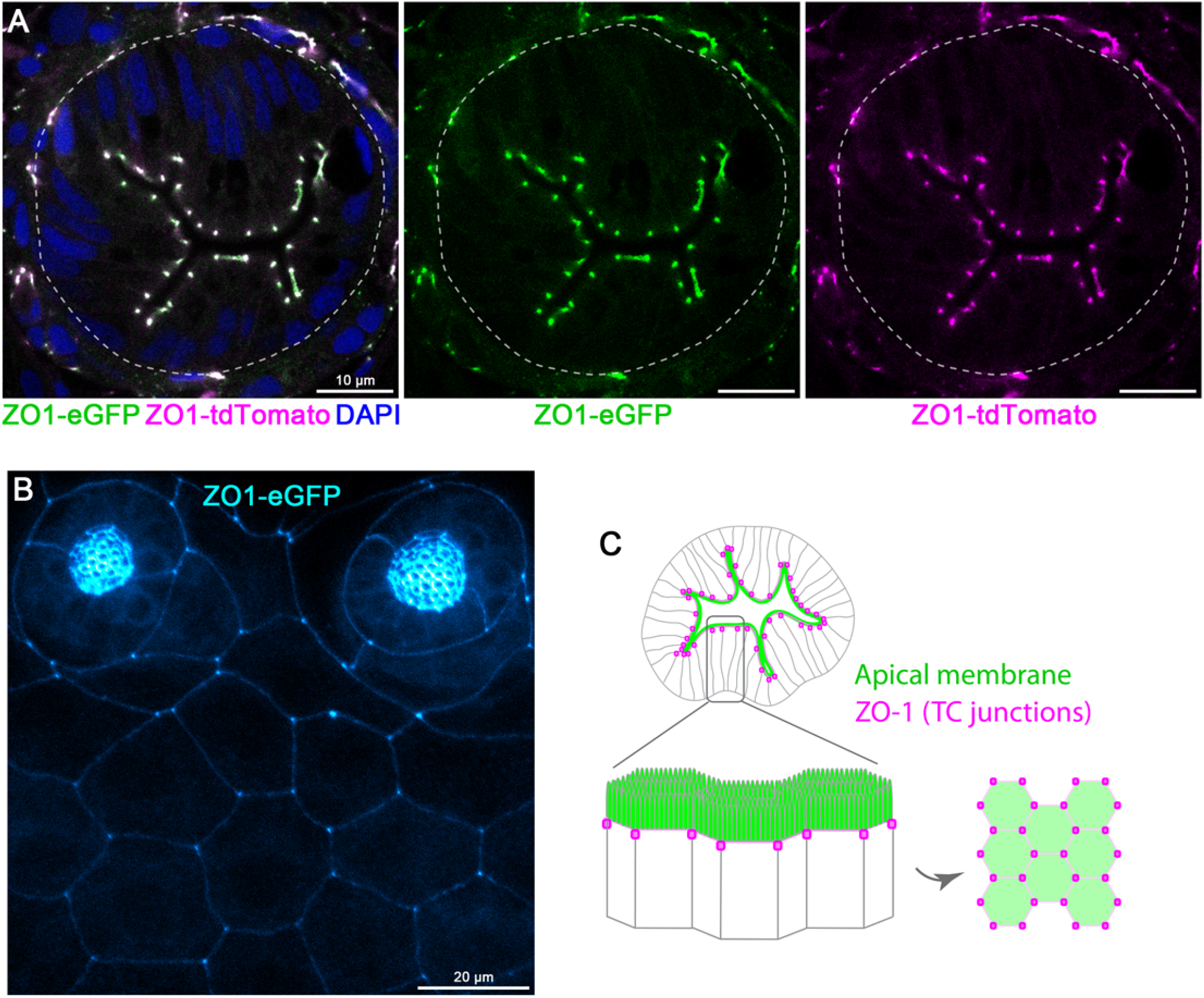
Colocalization of compound heterozygous tagged ZO1 (*tjp1a*) alleles and localization to tricellular junctions. **(A)** Transverse sections of a *TgKI(tjp1a-eGFP)*^pd1252^;*TgKI(tjp1a-tdTomato)*^pd1224^ compound heterozygous larva at 5 dpf. Scale bars are 10 μm. **(B)** Live imaging of the epidermis of a 5 dpf *TgKI(tjp1a-eGFP)*^pd1252^ larva. Panel B is pseudo-colored with the ImageJ/FIJI Cyan Hot LUT. Scale bars is 20 μm. **(C)** Schematic illustrating the enriched localization of endogenously tagged ZO1 to tricellular (TC) junctions.

